# M^6^A reduction relieves FUS-associated ALS granules

**DOI:** 10.1101/2023.10.25.563954

**Authors:** Gaia Di Timoteo, Andrea Giuliani, Adriano Setti, Martina C. Biagi, Michela Lisi, Alessia Grandioso, Davide Mariani, Francesco Castagnetti, Eleonora Perego, Sabrina Zappone, Giuseppe Vicidomini, Dante Rotili, Mario Sabatelli, Serena Lattante, Irene Bozzoni

**Author notes:** The authors wish it to be known that, in their opinion, the first two authors should be regarded as joint First Authors.

## Abstract

Amyotrophic lateral sclerosis (ALS) is a progressive neurodegenerative disease due to gradual motorneurons (MN) degeneration^1^. Among the processes associated to ALS pathogenesis, there is the formation of cytoplasmic inclusions produced by mutant protein aggregation, among which the RNA binding protein FUS^2^.

In this work we show that such inclusions are significantly reduced in number and dissolve faster when the RNA m^6^A content is diminished as a consequence of the m^6^A writer METTL3 knock-down. These effects were observed both in neuronal cell lines and in iPSC-derived human motor neurons expressing mutant FUS. Importantly, stress granules formed in ALS condition showed a distinctive transcriptome with respect to control cells; interestingly, after METTL3 downregulation, it reverted to similar to control. Finally, we show that FUS inclusions are reduced also in patient-derived fibroblasts treated with STM-2457, a well characterized inhibitor of METTL3 activity, paving the way for its possible use for counteracting aggregate formation in ALS.

## Introduction

Amyotrophic lateral sclerosis (ALS) is a progressive neurodegenerative disease. The gradual motoneurons degeneration caused by ALS leads to the atrophy of the innervating muscles and consequently to paralysis and death^1^. No cure for ALS exists today, with the only exception of a treatment specific for patients carrying pathogenic variants in *SOD1* gene^3^.

Many cellular processes have been associated to ALS pathogenesis. Among them the formation of aberrant stress granules (SG) due to mutant proteins which lead to liquid-solid phase-transition and protein aggregation^2^. Physiological SG are ribonucleoprotein membrane-less organelles that form to protect cells from stress conditions. They are composed of mRNAs, RNA-binding proteins, and translation initiation factors that are sequestered in a reversible manner to regulate mRNA stability and translation under stress conditions. SGs can be induced by a variety of stressors, such as oxidative stress, hypoxia, and heat shock, and are thought to represent a key adaptive response to cellular stress ^4^.

Usually, SG are transient structures that disassemble when the stress is over, but they can become pathological and cause cell death when stress is prolonged. Defects in both SG assembly and disassembly have been linked to neurodegenerative disorders^2,5,6^. Several mutant proteins associated to ALS, such as TDP-43 and FUS, localize, together with SG markers, in cytoplasmic inclusions commonly found in ALS patient samples^5–7^.

N6-methyladenosine (m^6^A) is a RNA modification known for its role in many biological processes^8^. However, very little is known about its role in the dynamics of physiological or pathological condensates.

Although m^6^A modification has been suggested to enhance the phase separation potential of mRNA *in vitro*^9,10^, it has been shown to be unable to play a significant role in mRNA recruitment in SGs *in vivo*^11^. In this study we investigated the interplay between SG and m^6^A in the context of ALS demonstrating that while m^6^A plays a limited role in SG physiology in control conditions, it has a strong impact in ALS genetic contexts, such as those where the FUS protein is mutated in domains that confer it a cytoplasmic relocation.

Indeed, using cellular models of ALS, we found that SG formed in the presence of mutant FUS differ from those of control cells not only in the number of condensates and in the rate of recovery from stress but also in RNA composition; Importantly, we show that the reduction of cellular m^6^A levels recovers these parameters towards those of control cells.

Interestingly, inhibition of METTL3 activity through the inhibitor STM-2457 led to an effective reduction of FUS-SG formation both in neuronal cell lines as well as in fibroblasts derived from ALS patients. Finally, we observed a weaker FUS confinement within condensates when m^6^A levels are lowered. Collectively, these data might foresee a potential application of such approach in those cases where protein aggregates formation has pathological implications.

## Results

### METTL3 downregulation restores the physiological RNA composition of stress granules in ALS cellular models

In order to unveil the impact of m^6^A on SG RNA composition in physiological and ALS-linked conditions, we used SK-N-BE cells carrying the doxycycline inducible overexpression of either wild type FUS (FUS^WT^) or mutant FUS (FUS^P525L^), and constitutively expressing the SG marker G3BP1 tagged with GFP. SK-N-BE is a human neuroblastoma cell line widely used to model neurodegenerative diseases^12^.

Upon doxycycline induction (“D^+^”), we obtained a three folds overexpression of FUS (Fig. S1a). As expected, FUS^P525L^, due to the mutation in its nuclear localization signal, is not efficiently imported in the nucleus and associates into SG upon stress induction^13^ (Fig. S1b). Indeed, in this condition, its signal almost completely colocalized with that of G3BP1 (Fig. S1c).

We treated FUS^WT^ or FUS^P525L^ cells with a control shRNA or with the combination of two shRNAs against METTL3, the main m^6^A writer acting on mRNAs, obtaining about 50% reduction of the protein (Fig. S1d, Fig. 1A). Furthermore, we evaluated the m^6^A reduction on mRNAs upon METTL3 decrease through the colorimetric EpiQuik assay and verified that the reduction was comparable with previously published works^14–16^ (20%, Fig. S1e). Upon doxycycline treatment and oxidative stress induction with sodium arsenite (“Ars^+”^), we isolated SG through immunoprecipitation with anti-GFP antibodies^17,18^ and we proceeded to RNA-seq (Fig. 1A, Supplementary Table 1).

**Fig 1.**
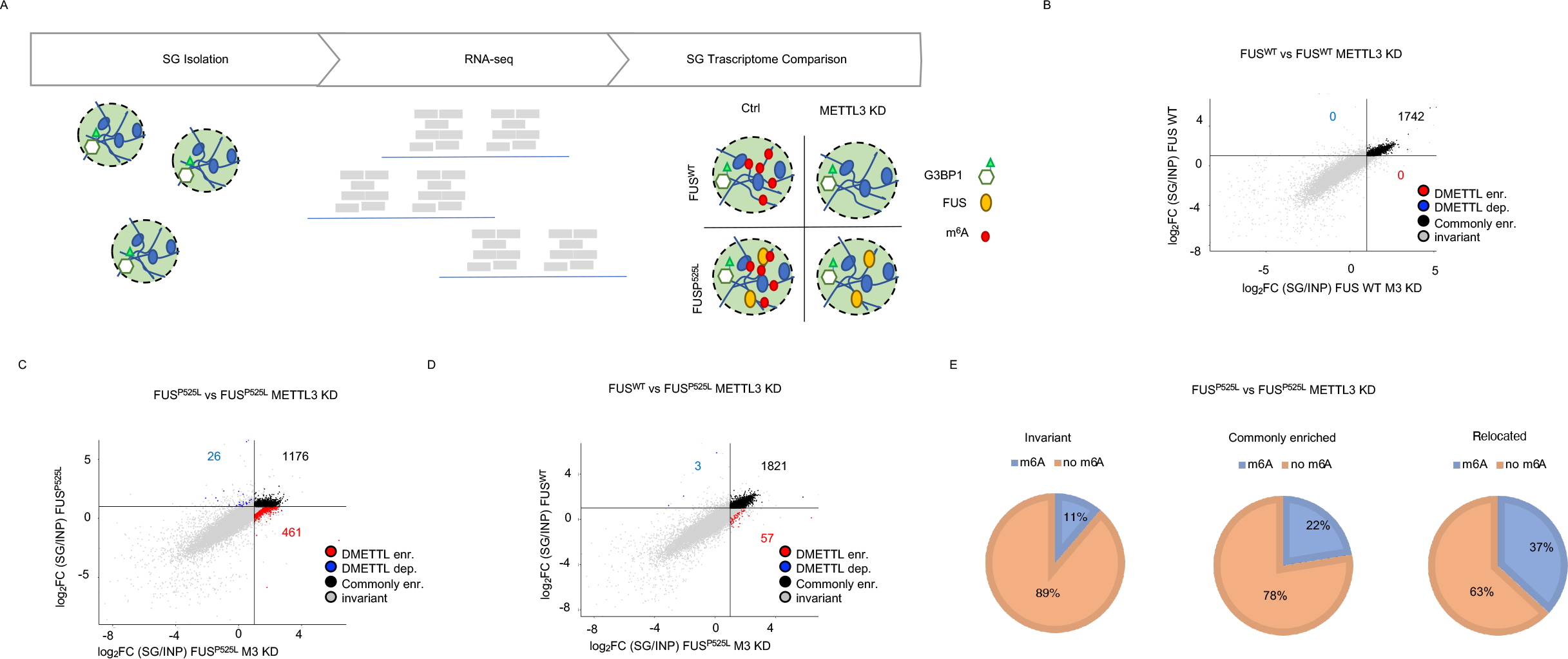
METTL3 downregulation restores the physiological RNA composition of stress granules in ALS cellular models. **a** Schematic representation of the experimental flow adopted for SG transcriptomes comparison. **b, c, d** Scatter plot depicting RNA differential enrichment in SG in FUS^WT^ vs FUS^WT^ METTL3-KD (b) or FUS^P525L^ vs FUS^P525L^ METTL3-KD (c) or FUS^wt^ vs FUS^P525L^ METTL3-KD (d) conditions. Axes describes log_2_FC of SG RNA enrichment in the indicated conditions. Red dots indicate ΔΔMETTL3-enriched RNAs. Blue dots indicate ΔΔMETTL3-depleted RNAs. Black dots indicate commonly enriched RNAs that are enriched in SG in both conditions. Grey dots indicate invariant RNAs. n=2. **e** Pie charts displaying the percentage of meRIP-Seq enriched RNAs among invariant, commonly enriched and relocated defined in the comparison between FUS^P525L^ vs FUS^P525L^ METTL3-KD. n=2.

SG enrichment in the immunoprecipitated fraction was confirmed by the enrichment of positive controls such as HUWE1, TRIO and ANHAK and the lack of ATP5O mRNA used as a negative control (Fig. S1f).

In order to identify transcripts possibly altered in their association with SG upon m^6^A reduction, RNA-seq data were analyzed through “Differential Enrichment Analysis” (DEA) comparing enrichment in SG in the presence of METTL3 and upon its downregulation. We distinguished four groups of transcripts: 1) RNAs significantly enriched in SG both in control and METTL3 KD conditions, defined as “*commonly enriched*”; 2) transcripts enriched only upon METTL3 depletion, named “*ΔMETTL3-enriched”;* 3) transcripts not enriched upon METTL3 depletion, defined as “*ΔMETTL3-depleted*”; 4) RNAs not enriched or showing no significant difference in SG enrichment, defined as “*invariant*”.

In agreement with previous data obtained in mES cells^19^, comparing the SG-associated transcriptome of FUS^WT^ cells in the presence or absence of METTL3, we did not observe any differentially enriched RNA, suggesting that m^6^A does not alter SG RNA composition in wild type conditions (Fig. 1b). No differences were also found in cells expressing only the endogenous FUS (Fig. S1g, “D^-^”), indicating that the increased levels of FUS expression did not affect SG composition. In contrast, as recently described^18^, we found strong differences when comparing SG composition of FUS^WT^ *versus* FUS^P525L^ condition, with 599 gained and 1267 lost transcripts in the mutant background (Fig. S1h). Remarkably, upon METTL3 downregulation, SG composition of FUS^P525L^ cells changed and returned very similar to that of FUS^WT^ relocating in SG a conspicuous fraction of RNAs (Fig. 1c, Fig. 1d). In fact, out of the 461 transcripts specifically recruited in FUS^P525L^-containing SG, ∼75% resulted in common with FUS^WT^ SG. For this reason, we will refer to such RNAs as “*relocated RNAs*” (Fig. S1h).

This evidence was also confirmed by the differential enrichment analysis performed by directly comparing FUS^WT^ *versus* FUS^P525L^ in conditions of METTL3 downregulation: in fact, we observed a strong similarity between these two SG transcriptomes, with only 57 differentially enriched transcripts (Fig. 1d). Indeed, while the FUS^P525L^ SG transcriptome displayed strong differences with FUS^WT^ or D-control conditions, upon shMETTL3 the SG of wild type and mutant FUS displayed similar RNA composition (Fig. S1i). Through enrichment convergence and resampling analyses, we verified that this similarity was not dependent on the threshold adopted to define enriched RNAs in SGs or on the RNA-Seq library size variability (Fig. S1j, Fig. S1k).

In agreement with these results, while the nucleotide composition of FUS^WT^ and FUS^P525L^ SG in control conditions was significantly different^18^, it returned similar following METTL3 downregulation (Fig. S1l).

Overall, these data indicate that while FUS^WT^ and FUS^P525L^ SG transcriptomes are markedly different, they revert similar under conditions of m^6^A downregulation.

In order to correlate the methylation *status* of the transcripts with their localization, we performed MeRIP-Seq in FUS^P525L^ cells (Supplementary table 2). The correct purification of methylated RNA was testified by the significant over-representation of the consensus DRACH motif in peaks regions (Fig. S1m). Furthermore, as expected^20^, the metagene plot resulting from these data displayed signal density near the stop codon (Fig. S1n). Finally, we validated by RT-qPCR the m^6^A content of several RNA species identified in the MeRIP-seq, confirming the correct immunoprecipitation of m^6^A-containing RNA (Fig. S1o).

When analyzing the *commonly enriched* RNAs in FUS^P525L^ SG, we found that m^6^A-containing RNAs accounted for 22% while those of *invariant* species corresponded to 11% (Fig. 1e). Such difference is mainly due to an increase in the length of the RNA (Fig. S1p), further confirming previous observations that long transcripts are enriched in SG and are more likely to contain m^6^A^17,19^.

Instead, the percentage of m^6^A-containing RNAs increased up to 37% when looking at the *relocated RNA* (Fig. 1e). Importantly, differently from the previous RNA subset, this enrichment is not dependent on the differential length of the transcripts since it resulted comparable to that of other transcripts included in the granules (Fig. S1p).

The fact that, in ALS-like conditions, the downregulation of METTL3 restored the wild type SG composition, together with the observation that there is a higher percentage of m^6^A-containining RNAs among the transcripts included in FUS^P525L^ SG, suggested a link between m^6^A modification and altered SG dynamics in ALS.

### METTL3 downregulation reduces the number of SG in ALS cellular models

In order to investigate if the downregulation of METTL3 could impact not only on the RNA content of FUS^P525L^ SG, but also on their number and size, we combined immunostaining for FUS and G3BP1 (Fig. 2a).

**Fig 2.**
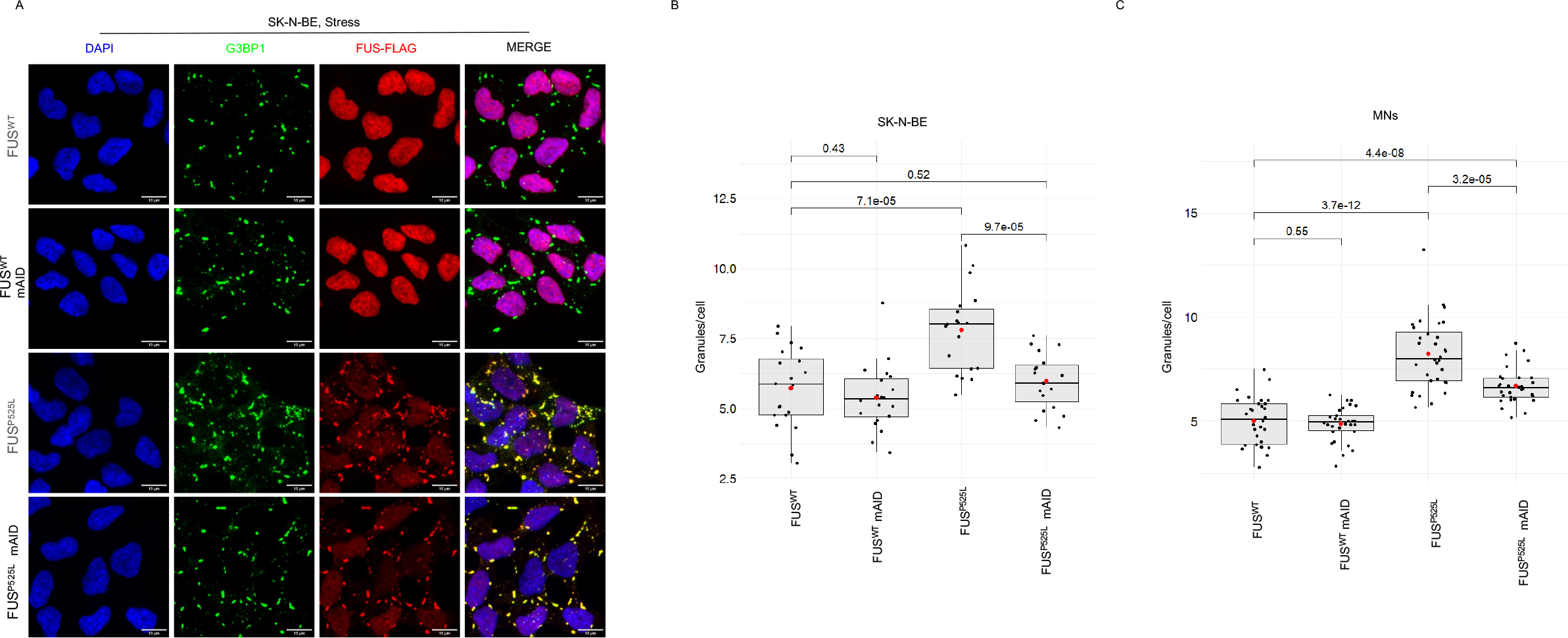
METTL3 downregulation reduces the number of SG in ALS cellular models. **a**, Representative images of the indicated stressed SK-N-BE cells. G3BP1 antibody staining is illustrated in green, FUS-Flag in red. The nuclei are stained with DAPI (blue). The merge of the signal is shown. The scale bar is 10 *μ*m. **b**,**c**, Box plots representing the number of stress granules per cell in the indicated SK-N-BE cell lines (n=3) or iPS-derived MN (n=4). The red dot indicates the average number of granules in each sample after 1hr stress. Each black dot represents the number of granules in a single field. Seven fields were acquired for each biological replicate. n=3.

Aiming to obtain stable and homogenous METTL3 knock-down conditions, we used the CRISPR-Cas9 technology to insert a degron tag (mAID) at the N-terminus of METTL3^21^ in order to obtain its auxin-inducible degradation (Fig. S2a). We obtained only heterozygous clones in both FUS^WT^ and FUS^P525L^ cell lines (Fig. S2b). These clones showed a decrease of METTL3 both at protein (∼50%, Fig. S2c, S2d) and RNA (∼40%, Fig. S2e) levels, even without auxin induction. Notably, the downregulation of METTL3 in these conditions produced a similar decrease of its interacting partner METTL14^22^ (Fig. S2c, Fig. S2d).

Furthermore, in order to check whether the level of METTL3 downregulation was sufficient to obtain variation in the amount of m^6^A mRNA, we applied the EpiQuik m^6^A RNA methylation quantification assay (∼20-30% reduction, Fig. S2f).

In consideration of these data and of the evidence that the expression levels of FUS^WT^ and FUS^P525L^ upon doxycycline induction was not impaired by prolonged METTL3 downregulation (Fig. S2g), we adopted the mAID-METTL3 cell lines as stable METTL3 knock-down systems for imaging studies. Furthermore, with this aim, we also confirmed that these cells, mirroring the shRNA conditions, had SG with a similar transcripts enrichment; in particular, selected candidates belonging to the “*relocated RNAs*” group resulted specifically less represented in FUS^P525L^ SG in the presence of physiological levels of m^6^A with respect to all the other conditions (Fig. S2h).

We then counted the number of SG per cell upon arsenite treatment in control and in mAID-METTL3 cells expressing either FUS^WT^ or FUS^P525L^. In the presence of physiological levels of m^6^A, we observed a conspicuous increase of SG in FUS^P525L^ cells compared to FUS^WT^ (Fig. 2b). When we analyzed mAID-METTL3 cells, we did not observe any significant variation in SG number in FUS^WT^ cells; instead, we noted a great reduction in FUS^P525L^ cells, where the number returned to levels comparable to those of FUS^WT^ (Fig. 2b). Evaluation of SG volumes revealed no relevant average variation across the different conditions (Fig. S2i). However, when granules were divided by size into small (<0.1 *μ*m^3^), medium (0.1 *μ*m^3^-1 *μ*m^3^) and large (>1 *μ*m^3^), we noticed that the number of large granules per cell was maintained across the samples examined, while the number of small and medium granules were responsible for their statistically significant difference (Fig. S2j).

Since ALS affects MN, we performed the same analysis in human iPSC-derived MN expressing endogenous levels of FUS^WT^ or FUS^P525L^ ^23^.

In these cells we were able to obtain homozygous clones for the insertion of the degron-tag at the N-terminus of METTL3 (Fig. S2k), leading to an almost complete depletion of the protein, even in the absence of auxin, that was also paralleled by a strong downregulation of METTL14 (Fig. S2l). iPSCs were converted into MN according to Garone *et al*.^24^(2019) and analyzed after eleven days of trans-differentiation. Immunofluorescence experiments for METTL3 confirmed its decrease in these iPSC-derived MNs (Fig. S2m). Moreover, we also verified m^6^A RNA methylation decrease in mAID-METTL3 MN thanks to the EpiQuik quantification assay, which indicated an approximate ∼30-40% reduction of RNA m^6^A levels (Fig. S2n). The correct trans-differentiation was testified by the proper expression of three of the main MN markers (TUJ1, CHAT and ISLE-1), that was unaltered even upon METTL3 downregulation (Fig. S2o).

We then compared the number and volumes of SG in FUS^WT^ or FUS^P525L^ MN, with or without METTL3 depletion (Fig. 2c, Fig. S2p, Fig. S2q). The results confirmed what observed in SK-N-BE cells, namely that FUS^P525L^ cells produce a higher number of SG which are recovered to normal levels upon METTL3 downregulation with no relevant average variation in their volumes (Fig. 2c, Fig. S2q). In this case, when dividing SG based on their volumes, we could observe changes in both small (<0.1 *μ*m^3^), medium (0.1 *μ*m^3^-1 *μ*m^3^) and large (>1 *μ*m^3^) granules (Fig. S2r).

In conclusion, although METTL3 decrease did not affect SG number and size in a physiological context where wild type FUS is expressed, it reduced the number of SG to levels similar to the control conditions in the FUS-associated ALS models tested. Furthermore, these data indicate that the effect of m^6^A on SG dynamics is the same in cells expressing either the endogenous or the overexpressed mutant FUS.

### METTL3 decrease restores SG recovery rate in ALS cellular models

SG usually dissolve when stress is removed and defects in such process are linked to several human neurodegenerative diseases ^25–27^.

In order to investigate the effects of m^6^A on SG disassembly, we evaluated the recovery rate in METTL3 knock-down conditions through immunofluorescence analysis (Fig. 3a).

**Fig 3.**
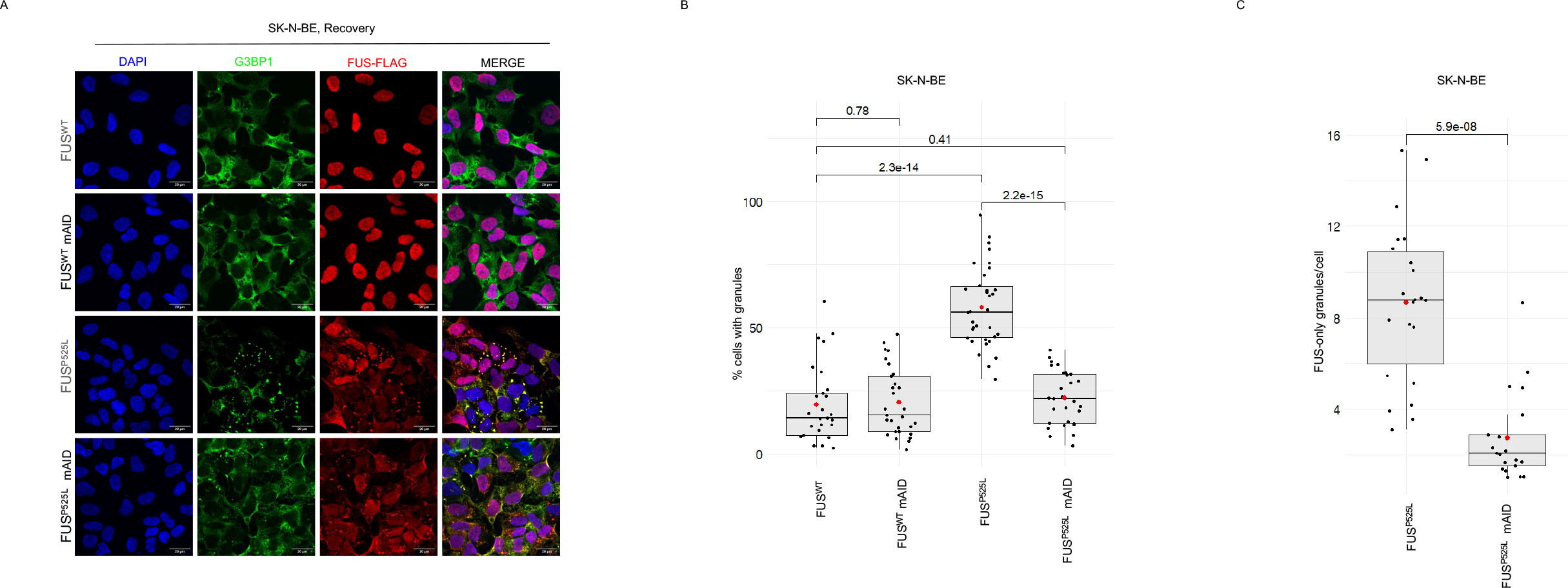
METTL3 decrease restores SG recovery rate in ALS cellular models. **a**, Representative images of the indicated SK-N-BE cells after 1hr stress followed by 4hr recovery. G3BP1 antibody staining is illustrated in green, FUS-Flag in red. The nuclei are stained with DAPI (blue). The merge of the signal is shown. The scale bar is 20 *μ*m. **b**, Box plots showing the percentage of unrecovered indicated SK-N-BE cells after 1hr stress followed by 4hr recovery. The red dot indicates the average percentage in each sample. Each black dot represents the percentage in a single field. Seven fields were acquired for each biological replicate. n=4 biologically independent replicates. The *ratio* of each sample versus its experimental control was tested by two-tailed Student’s t test. P-values are indicated. **c**, Box plots showing the number of FUS-only granules in the indicated FUS^P525L^ SK-N-BE cells. The red dot indicates the average granules number in each sample. Each dot represents the number of granules in a single field. Seven fields were acquired for each biological replicate. n=4 biologically independent replicates. The *ratio* of each sample versus its experimental control was tested by two-tailed Student’s t test. P-values are indicated.

Fig. 3b shows the percentage of FUS^WT^ and FUS^P525L^ SK-N-BE cells still containing granules after 4hr recovery from stress removal, either in control conditions or upon METTL3 downregulation. The results indicate that FUS^P525L^ cells showed a higher percentage of unrecovered cells with respect to FUS^WT^ (Fig. 3b). Moreover, while the depletion of the m^6^A writer did not change the recovery rate of FUS^WT^, in mutant cells it rescued the recovery rate to wild type levels (Fig. 3b).

Interestingly, while we observed almost total overlap of G3BP1 and FUS signals upon stress in FUS^P525L^ cells (Fig. S1c), we noticed granules displaying FUS-only signal after recovery (Fig. S3a). Coherently, colocalization analysis showed that the observed total overlap of the two signals under stress conditions was significantly reduced to about 40% under recovery condition (Fig. S3b). Notably, the number of FUS-only aggregates decreased upon m^6^A downregulation (Fig. 3c), and these condensates resulted smaller than the granules observed in stress conditions (Fig. S3c).

These observations suggest the hypothesis that, upon stress, SG containing both G3BP1 and FUS^P525L^ are formed but that, after stress removal, G3BP1 is released free into the cytoplasm while FUS persists in an aggregate form. More importantly, FUS aggregation is relieved if the levels of m^6^A are reduced.

### METTL3 chemical inhibition relieves FUS-containing SG

Given the availability of a small molecule, STM-2457, which has been previously validated as an effective and specific inhibitor of METTL3^28^, it was worth it to verify whether it could reproduce some of the phenotypes observed in ALS cellular models.

We initially tested its activity in SK-N-BE cells expressing FUS^P525L^. Figure S4a shows the effective m^6^A decrease upon STM-2457 treatment (15μM) with respect to the DMSO control (∼75% m^6^A reduction).

The decrease of m^6^A levels due to STM-2457 administration recapitulated the effects of the lack of METTL3: indeed, cells formed less granules and displayed a higher recovery rate (Fig. S4b, Fig. S4c).

Given the availability of patient-derived fibroblasts carrying the mutant FUS^R518I^ allele, we applied the STM-2457 treatment to both these cells and iPSC-derived MNs carrying FUS^P525L^. The FUS^R518I^ mutation, although different from the highly pathogenic FUS^P525L^, also causes relocalization of FUS in the cytoplasm, albeit to a lower extent, and causes the formation of FUS-containing SG upon stress induction (Fig. 4a).

**Fig 4.**
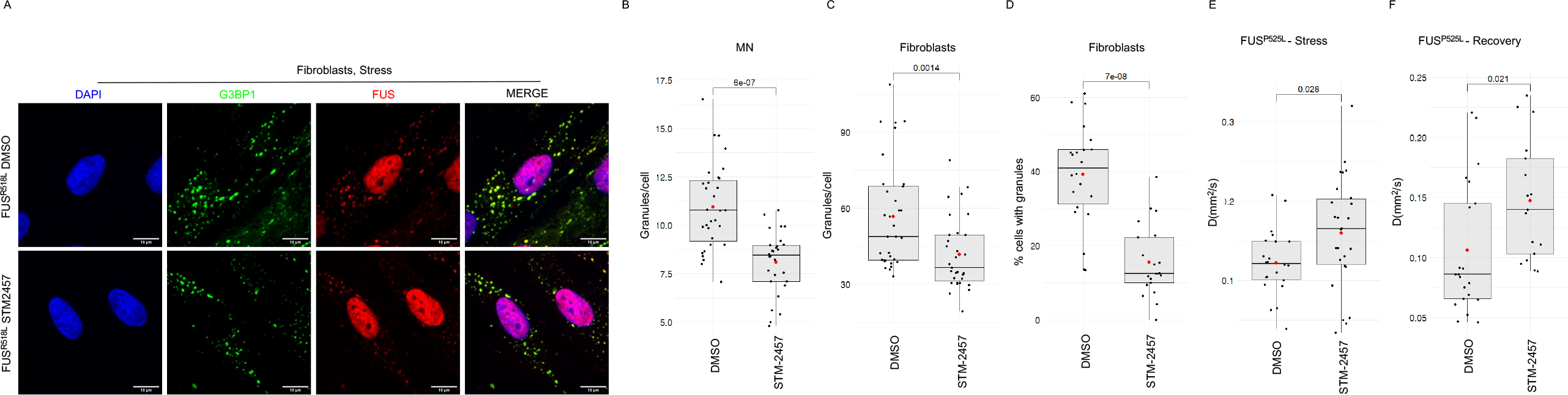
METTL3 chemical inhibition relieves FUS-containing SG. **a**, Representative images of the indicated stressed patients-derived fibroblasts. G3BP1 antibody staining is illustrated in green, FUS in red. The nuclei are stained with DAPI (blue). The merge of the signal is shown. The scale bar is 10 *μ*m. **b**,**c**, Box plots representing the number of stress granules per cell in stressed iPS-derived MN expressing FUS^P525L^ or patient-derived fibroblasts treated either with DMSO or STM-2457. The red dot indicates the average number of granules in each sample. Each black dot represents the number of granules in a single field. Seven fields were acquired for each biological replicate. n=4 biologically independent replicates. The *ratio* of each sample versus its experimental control was tested by two-tailed Student’s t test. P-values are indicated. **d**, Box plots showing the percentage of unrecovered patients-derived fibroblasts treated either with DMSO or STM-2457. The red dot indicates the average percentage in each sample. Each black dot represents the percentage in a single field. Seven fields were acquired for each biological replicate. n=3 biologically independent replicates. The *ratio* of each sample versus its experimental control was tested by two-tailed Student’s t test. P-values are indicated. **e**,**f**, Box plots representing the diffusion coefficients (D) of FUS^P525L^ in living SK-N-BE cells treated with either DMSO or STM-2457 in stress or recovery condition. The red dot indicates the average diffusion coefficient value. Each black dot a single measurement. n=2 biologically independent replicates. The *ratio* of each sample versus its experimental control was tested by two-tailed Student’s t test. P-values are indicated.

For these cellular models we assessed the m^6^A decrease on RNAs with the EpiQuik assay showing its efficient decrease (∼70-80%) of m^6^A deposition (Fig. S4d and S4e).

Figures 4b and 4c show how the inhibition of METTL3 activity significantly reduces the number of stress granules in iPSC-derived MNs as well as in patient-derived fibroblasts. Furthermore, in line with our previous results, we observed a lower percentage of cells with granules in patient-derived fibroblasts recovering from stress when treated with STM-2457 (Fig. 4d).

We also employed a variation of conventional fluorescence correlation spectroscopy, utilizing a single-photon avalanche diode (SPAD) array detector^29,30^. This innovative approach enables comprehensive studies of molecular mobility within living cells. In particular, we measured FUS^P525L^ and G3BP1 diffusion coefficient (*D*) and their confinement strength (*S*_conf_) both under stress and recovery, either in control conditions or upon METTL3 inhibition in SK-N-BE cells expressing the GFP-tagged versions of FUS or G3BP1 (Fig. 4e, Fig. S4f-k).

The diffusion coefficient is related to the movement of the molecules: the higher the value of *D*, the faster is the diffusion rate of the molecule. The confinement strength is a metric related to the type of movement: *S*_conf_ values below one correlate with strong molecule confinement, while values closer to one indicate a free diffusion behavior.

G3BP1 did not undergo noteworthy *S*_conf_ or *D* alterations upon METTL3 inhibition both in stress and recovery conditions, meaning that its movement is not affected by m^6^A reduction (Fig. S4h, Fig. S4i, Fig. S4j, Fig. S4k). Coherently with our data, METTL3 inhibition provoked instead a significant increase of both *D* and *S*_conf_ values of FUS^P525L^, indicating a higher diffusion coefficient and a lower confinement with respect to the control condition (Fig. 4e, Fig. 4f, Fig. S4f, Fig. S4g). These data confirm that the reduction of m^6^A levels relieves aggregation allowing FUS^P525L^ to be involved in less dense structures.

## Discussion

RNA-protein (RNP) assemblies vary in a dynamic manner^6^ and changes in the relative abundance of proteins and RNAs can lead to liquid-solid phase transition and eventually to the formation of aggregates^31,32^. RNP assemblies acquire significant relevance in neurodegenerative diseases where protein mutations and physicochemical insults can induce liquid-solid phase-transition^31^.

ALS is associated with appearance of protein aggregates in MNs^33^ and several findings causatively link this phenomenon with pathological states: in fact, overexpression of either wild-type or mutant RBPs such as FUS, TDP-43 (in ALS and Frontotemporal disorders) and FMRP (in Fragile X Tremor Ataxia Syndrome) in different model organisms resulted in reduced motility and processivity of mitochondrial axonal transport and altered displacements of RNP assemblies^34–36^. Spatial localization of RNA is a very crucial process in neurons, as it controls local protein synthesis that mediates directional and morphological responses. Transport defects have been ascribed to RBPs and their ability to control correct subcellular compartmentalization of several macromolecular structures, from large organelles to mRNAs and their localized translation^37^.

Therefore, the ability to prevent or revert aggregate formation could represent an important therapeutic approach for several neurodegenerative pathologies. As proof of this, molecular chaperones and pharmacological induction of autophagy have been shown to decrease FUS aggregation and to alleviate associated cytotoxicity, resulting in improved neuronal survival and function^38,39^. Furthermore, antisense oligonucleotides targeting FUS have been shown to decrease FUS aggregation and rescue RNA processing defects in cellular models^40^.

In this work, we demonstrate a new approach to reduce the formation of mutant FUS-containing SG and to facilitate their dissolution, consisting in reducing the levels of m^6^A deposition on RNA. Interestingly, reduction of FUS-containing SG condensates was obtained in cell lines and in patient-derived fibroblasts also with the small molecule STM-2457, a known fully validated inhibitor of the methyltransferase activity of the METTL3-METTL14 complex.

Notably, the phenotypes observed in the presence or depletion of m^6^A were paralleled by an interesting reshaping of the transcriptomes: while the RNA content of SG obtained in FUS^P525L^ was considerably different from that of FUS^WT^, when m^6^A was reduced, the transcriptome reverted towards the control one. These data indicated the important role of m^6^A in contributing to the phase transition occurring in SG of FUS^P525L^ cells, suggesting that the modulation of m^6^A plays a relevant role in aberrant SG formation and indicating that targeting this modification can indeed revert not only the morphology but also the RNA composition of these condensates.

The specific molecular mechanism underlying RNA relocation is yet to be understood; further work will be required to investigate the link between m^6^A and FUS RNA binding properties and how they are affected by FUS mis-localization in the cytoplasm and by its potentially aberrant novel interactions. Nevertheless, our data indicate that physiological levels of m^6^A favor condensation in FUS^P525L^ conditions while its reduction promotes SG fluidification.

Finally, the finding that STM-2457 phenocopies the decrease of FUS aggregates in METTL3 knock-down cell lines opens the perspective of repurposing this molecule, previously adopted in clinical trials as an inhibitor of cell proliferation in the therapeutic treatment of acute myeloid leukemia^28^, for the inhibition of protein aggregates in FUS-associated ALS cases.

Further studies will allow to test whether this type of treatment can be effective also for counteracting other neurodegenerative diseases causally linked to the accumulation of pathological protein aggregates, such as Alzheimer and Parkinson diseases.

## Author Contributions

G.D.T., A.G. and I.B. designed and conceived the study. The experiments were performed and analyzed by G.D.T., A.G., M.C.B., M.L., A.G., D.M., F.C., E.P., S.Z., G.V., D.R., M.S., S.L..

Bioinformatics data analysis was performed by A.S. The original draft of the manuscript was written by I.B., G.D.T. and A.G. with suggestions from all the other authors. I.B. supervised the project.

## Data Availability

All software, links to websites or tools used for this work are referred to in the Materials and Methods section or in the figure legends. Additional dedicated scripts developed for this work are available upon request.

## Inclusion & Ethics statement

Human primary fibroblasts were obtained from a patient carrying the p.R518I in *FUS* gene, after written informed consent and approval of the Ethic Committee (Protocol nr. A.133/CE/2013).

## Code availability statement

All software and tools used for this work are described in the “Methods” section or in the figure legends. Additional dedicated scripts developed for this work are available from the corresponding authors upon request.

## Acknowledgements

We thank M. Morlando, A. Fatica, V. De Turris, A. Rosa, J. Martone, T. Santini, J. Rea, A. Colantoni for useful discussions and suggestions. We also thank F. Margarita and M. Marchioni for technical help and Dr M. Caruso for assistance.

This work was supported by grants from ERC-2019-SyG 855923-ASTRA, AIRC IG 2019 Id. 23053, PRIN 2017 2017P352Z4 to I.B.; “National Center for Gene Therapy and Drugbased on RNA Technology” (CN00000041), NextGenerationEU PNRR MUR to I.B. and G.V.; ERC-2018-CoG 818669-BrightEyes to G.V.; “Sapienza” Ateneo Project 2021 n. RM12117A61C811CE and Regione Lazio PROGETTI DI GRUPPI DI RICERCA 2020 - A0375-2020-36597, NextGenerationEU through the Italian Ministry of University and Research under PNRR - M4C2-I1.3 Project PE_00000019 “HEAL ITALIA” CUP (B53C22004000006) to D.R..

## Methods

### Cell culture and preparation

SK-N-BE cells were cultured in RPMI (Sigma-Aldrich, Saint Louis, MO, USA) supplemented with 10% FBS (Sigma-Aldrich, #F2442), GlutaMAX supplement 1X (ThermoFisher Scientific, # 35050061), sodium-pyruvate 1mM (Thermo Fisher Scientific, #11360070) and Pen/Strep 1X (Sigma-Aldrich, #P4458). For the inducible expression of FUS-FLAG WT or P525L, SK-N-BE cells were exposed to 50 ng/mL Doxycycline (Sigma-Aldrich, #D9891) for 24h before the sodium arsenite treatment (Sigma-Aldrich, #106277).

For the immunofluorescence experiments, SK-N-BE cells were dissociated with Trypsin 1X (Sigma-Aldrich, #T4549) and 20,000-30,000 cells/well were plated on glass coverslips in 24-well plates.

Skin biopsy was performed at the distal leg using a 4-mm punch. Skin samples were dissected and cultured in BIOAMF-2 (Biological industries) complete medium. Human primary fibroblasts were cultured in DMEM high-glucose with 1mM sodium-pyruvate (Sigma-Aldrich, # P5280) supplemented with 20% FBS, L-glutamine 2mM and Pen/Strep 1X. Cells were dissociated with trypsin-EDTA 1X (Corning, #25-053-CI) and 50,000 cells/well were plated on collagen-coated glass coverslips in 24-well plates. The following day, fibroblasts were treated with 15*μ*M STM-2457, while control fibroblast were treated with same volumes of DMSO (Sigma Aldrich, #D2650), for 48h before the sodium arsenite treatment.

Human NIL iPSCs were cultured in Nutristem (Sartorius, #05-100-1A) supplemented with Pen/Strep 0.1% on geltrex-coated plates (Thermo Fisher Scientific, #A1413202) and differentiated towards spinal motoneurons as described by Garone et al. 2019^24^. After 5 days of differentiation, cells were dissociated with Accutase (Thermo Fisher Scientific, #00-4555-56) and 200,000 cells/well were plated on geltrex-coated glass coverslips in 24-well plates. After additional 6 days of differentiation, the iPSC-derived motoneurons were treated with sodium arsenite. As regards for METTL3-inhibition experiments, MNs were treated with 15μM STM-2457, while control MN were treated with same volumes of DMSO, for 48h before the arsenite treatment.

All cell lines used in were grown at 37°C, 5% CO2. All cell lines were tested for mycoplasma contamination.

For acute oxidative stress treatment, SK-N-BE cells, iPSC-MNs and primary fibroblasts were exposed to 0.5 mM sodium arsenite for 1 hr. Rescue experiments were performed by restoring normal growth medium for 4h or 2h after acute arsenite treatment in SK-N-BE cells or fibroblasts, respectively.

### Chemicals

STM-2457 has been prepared according to published procedures^41^.

### Genome editing

mAID-tag was inserted at the N-terminus of METTL3 together with the coding sequence for the hygromicyn resistance and a p2A signal for a proteolytic cut in SK-N-BE and iPS cells by Crispr/Cas9-induced DNA break. Two guide RNAs targeting the 5’UTR region of METTL3 were designed with CHOPCHOP (https://chopchop.cbu.uib.no/) and cloned in a px330 vector through BbsI digestion. The plasmid donor carrying the sequence to be inserted plus 500nt of homology upstream and downstream was synthetized by GENEWIZ. 10^6^ SK-N-BE cells were transfected with 3*μ*L of Lipofectamine 2000 (Life technologies) in 300μL of Optimem with donor 2 ug of DNA donor and 2 ug of each guide plasmid. Same amount of DNA was transfected in 10^6^ iPS cells with Neon™ Transfection System 100*μ*L Kit (Invitrogen, #MPK10096) according to manufacturer’s instruction (1 pulse, 20 V, 30ms). Cells transfected with donor only were used as selection control. Medium was replaced with fresh one added with 450*μ*g/mL or 300*μ*g/mL hygromycin 24hr after transfection for SK-N-BE and iPS cells, respectively. Selection was carried out until control cells were died. Single colonies were transferred to 24-well plates. Colonies were then split, and a half was used for gDNA extraction and genotyping through Rapid Extraction PCR kit (PCR biosystems, #PB10.24). Oligos 9F and 4R were used for amplifying the WT and recombinant alleles (Supplementary Table 2).

### Immunocytochemistry

Cells were fixed for 20min at RT with cold 4% paraformaldehyde (Electron Microscopy Sciences, #15710) diluted to in complete PBS (Sigma Aldrich, # D1283), rinsed 3 times with complete PBS and stored in PBS at 4°C. Cells were permeabilized with 0.3% Triton X-100 (Sigma-Aldrich, #648466) diluted in complete PBS for 10 min and blocked with 5% goat and/or donkey serum, depending on the host species of the secondary antibody used. Samples were then incubated overnight at 4°C with the primary antibody diluted in blocking solution (5% goat and/or donkey serum in PBS, #G9023 and #D9663 Sigma-Aldrich). Cells were washed with complete PBS 3 times for 5min at RT and then incubated with the following secondary antibodies for 1h at RT. diluted in blocking solution and were incubated for 1h at RT. Nuclei were stained with 1*μ*g/ml DAPI (#D9542, Sigma-Aldrich) diluted in complete PBS for 5 min and coverslips were mounted applying ProLong™ Glass Antifade Mountant (P36980, Thermo Fischer Scientific) leaving the slides on the bench. Confocal images were acquired with an inverted Olympus iX73 equipped with an X-Light V3 spinning disc head (Crest Optics), a Prime BSI Scientific CMOS (sCMOS) camera (Photometrics) and MetaMorph software (Molecular Devices), as Z-stacks (0.3 um step size) with a 60× oil-immersion objective.

The following antibodies were used for immunohistochemistry in this study:

a rabbit monoclonal anti-G3BP1 antibody (1:300, ab181150 Abcam)

mouse anti-FLAG antibody (1:400, F1804 Sigma-Aldrich)

mouse anti-FUS antibody (1:300, sc-47711 Santa Cruz Biotechnology)

rabbit monoclonal anti-METTL3 antibody (1:500, ab195352 Abcam)

chicken Anti-Beta III Tubulin (TUJ1) Antibody (1:400, AB9354 Sigma-Aldrich)

goat anti-rabbit Alexa Fluor 488 (1:300, A11008 Thermo Fisher Scientific),

goat anti-mouse Alexa Fluor 488 (1:300, A11001 Thermo Fisher Scientific)

goat anti-rabbit Alexa Fluor 555 (1:300, 111-165-003 Jackson Immunoresearch)

goat anti-mouse Alexa Fluor 555 (1:300, 115-165-003 Jackson Immunoresearch)

goat anti-chicken Alexa Fluor 594 (1:300, ab150176 Abcam)

donkey anti-mouse Alexa Fluor Plus 647 (1:300, A32787 Thermo Fisher Scientific),

donkey anti-rabbit Alexa Fluor Plus 647 (1:300, A32795 Thermo Fisher Scientific)

### Image Analysis

The analysis of immunofluorescence images was performed through the ImageJ software^42^. G3BP1 was used as SG marker. The ImageJ tool “3D Object Counter” was used for the quantification of SGs number and volumes. For the estimation of cells forming SGs in the recovery experiments cells were scored as SG-positive when they presented at least one G3BP1 granule-like cytoplasmic signal. For the intensity fluorescence analysis, a nuclear mask was generated using the DAPI signal to specifically select nuclear regions; this mask was then used to measure the METTL3 fluorescence intensity mean of each nucleus with the ImageJ tool “Measure”. For the signal colocalization analysis the ImageJ plugin “Jacop” was used to calculate the Manders coefficient of FUS signal colocalizing with G3BP1 signal. In order to consider only the cytoplasmatic signal of FUS, a nuclear mask was created for each image and removed from both the FUS and G3BP1 images prior to proceed with the analysis. For the signal profile analysis random granules were selected and sectioned with a 20-pixel length straight line and the ImageJ tool “Signal Profile” was used to quantify the signal intensity. For the quantification of “FUS-only” granules in recovery experiments, the G3BP1 signal was used to create a mask and then removed from the FUS signal before the quantification of the number and volumes of “FUS-only” granules using the ImageJ tool “3D Object Counter”.

### Protein analyses

Protein analyses were carried out as described in Dattilo et al. 2023^43^.

The following antibodies were used in western blots for protein analyses:

mouse anti-FUS antibody (1:300, sc-47711 Santa Cruz Biotechnology)

rabbit monoclonal anti-METTL3 antibody (1:500, ab195352 Abcam)

Anti-METTL3 [EPR18810] monoclonal antibody Abcam ab195352 1:1000

Anti-METTL14 polyclonal antibody Atlas HPA038002 1:1000

Anti-Flag M2-Peroxidase (HRP) Sigma-Aldrich A8592 1:2500

Anti-ACTB-Peroxidase (AC-15) monoclonal antibody Sigma-Aldrich A3854 1:2500

Anti-GAPDH (6C5) monoclonal antibody Santa Cruz Biotechnology sc-32233 1:1000

Anti-Rabbit IgG (H+L) Secondary Antibody, HRP Thermo Fisher Scientific 31460 1:10000

Anti-Mouse IgG (H+L) Secondary Antibody, HRP Thermo Fisher Scientific 32430 1:10000

### RNA analyses

RNA analyses were carried out as described in Dattilo et al.^43^.

### m^6^A RNA Methylation Quantification

After polyA+ RNA selection through Poly(A)Purist-MAG kit (Invitrogen, #AM1922), m^6^A level quantification was performed thanks to the EpiQuik m^6^A RNA Methylation Quantification Kit (EPIGENTEK, # P-9005) according to manufacturer’s instructions using 50-200 ng of RNA per well.

### Fluorescence correlation spectroscopy measurements

To perform the FCS measurements, we used a custom laser-scanning microscope equipped with a 5x5 SPAD array detector described by Perego et al.^30^ .

Cells were seeded onto a μ-Slide 8 well plate (Ibidi GmbH) and imaged in Live-Cell Imaging Solution (ThermoFisher Scientific) at 37°C. Before each measurement, the cells were visually inspected by imaging. The axial position for the spectroscopy measurements was placed in the middle of the chosen cell. Multiple planar positions in cells were selected to probe different points (in the cytoplasm or inside SGs). The fluorescence intensity was acquired for about 100 seconds and analyzed offline. All measurements were performed at 37°C inside a temperature-controlled chamber (Bold Line Temperature Controller, Okolab, PA, USA).

We calculated the time correlations directly on the lists of absolute photon times^44^). The data was then split into chunks of 10 s, and the time autocorrelations were calculated for each chunk. For the Sum 5×5 analysis, the lists of all SPAD channels were merged and the correlations were calculated. The individual correlation curves were visually inspected, and all curves without artifacts were averaged and fitted with a function describing a 3D diffusion. The diffusion coefficients are calculated from the fitted diffusion times knowing the confocal volumes of the microscope (calibrated as described by Perego et al. ^30^). Once the diffusion coefficients were calculated, the confinement strength is simply quantified by dividing the diffusion coefficient of the correlation of the signal from the central pixel only with the one retrieved from the correlation of the Sum 5×5 pixels.

### Stress granules isolation

2x10^6^ SK-N-BE cells FUS^WT^ or FUS^P525L^ were plated on a 6cm plate and transfected with 4*μ*g of vectors expressing sh1 and 4*μ*g of sh2 (TRCN0000289812, TRCN0000289814). The day after cells were transferred to a 15cm plates. When required, 96hr after transfection FUS expression was induced with doxycycline for 24hr, stressed 1hr before harvesting, and harvested according to the protocol described by Khong *et al*.^45^ In order to normalize on the immunoprecipitation efficiency, we used the invariant transcript CALM1 as endogenous control in the comparison of different SG isolation experiments.

### SG RNA-Seq analysis

Purified SG RNAs and relative inputs were generated in D-(4 replicates); upon FUS^P525L^ overexpression of (2 replicates) and FUS^WT^ (2 replicates) in condition of METTL3 depletion. RNA libraries for all samples were produced using Stranded Total RNA Prep with Ribo-Zero Plus (Illumina). All samples were sequenced on an Illumina Novaseq 6000 Sequencing system. Trimmomatic (v0.39) ^46^ and Cutadapt (v3.2) ^47^ were used to remove adapter sequences and poor-quality bases; minimum read length after trimming was set to 35. Reads aligning to rRNAs were filtered out; this first alignment was performed using Bowtie2 software (v2.4.2) (https://bowtie-bio.sourceforge.net/bowtie2/index.shtml). STAR software (v2.7.7a) ^48^ was used to align reads to GRCh38 genome using ENCODE standard parameters referred in the manual. PCR duplicates were removed from all samples using MarkDuplicates command from Picard suite (v2.24.1) (https://broadinstitute.github.io/picard/). Uniquely mapping fragments were counted for each annotated gene (Ensembl release 99) using HTseq software (v0.13.5) ^49^. EdgeR R package (v3.34.1) ^50^ was used to compare SG enriched RNAs to their relative input samples. RNAs with log_2_FC>1 and FDR<0.05 were defined “*enriched*”; those with log_2_FC < -1 and FDR < 0.05 were defined as “*depleted*”; all the other were labelled as “*invariant*”.

Differential enrichment analysis was performed using edgeR software defining the contrasts: *(IP1-INP1)-(IP2-INP2)* for the compared conditions. Significance threshold was set to FDR<0.05. Moreover, only RNAs expressed more than 1FPKM in at least one sample type (IP1 or INP1 or IP2 or INP2) and that resulted as SG “enriched” in one of the analyzed conditions were defined as differentially enriched. Differential enrichment was taken in consideration in order to define the “*ΔMETTL3-enriched”* and “*ΔMETTL3-depleted*” groups. In all the analyses that use transcript sequences, fasta files were retrieved from Ensembl using biomart ^51^. In case of genes with multiple isoforms a representative isoform was selected (the longest isoform with the same biotype of the gene was selected as representative; the longest isoform for non-coding RNAs).

In order to overcome the high variability related to immunoprecipitation-based experiments, we performed an Enrichment convergence analysis. For each analyzed condition, we first ranked the expressed RNAs by their SG enrichment (log_2_FC) and then we selected a fixed number of enriched RNAs for each dataset, and compared these three equally sized RNA sets. We repeated this analysis selecting different fixed numbers of enriched RNAs gradually reducing the inclusion of the noise from the background transcriptome (fixed numbers of top enriched RNAs: 5000, 4500, 4000, 3500, 3000, 2500, 2000, 1500, 1000, 500).

To avoid the technical bias due to the high variability of the library size of IP samples, we performed Resampling analysis in different conditions. We randomly sampled fragments from the alignment files of the other conditions in order to mirror the library size of the FUS^P525L^ condition experiment, used as reference and we repeated enrichment analyses. Random sampling of fragments was performed with Picard suite using FilterSamReads function and using random subsets of fragments ids list as input (https://broadinstitute.github.io/picard).

### K-mers analysis

The average transcript frequency of each k-mer was calculated for SG enriched RNAs and background distribution (invariant group), and their ratio (fold-change, FC) was computed. Then, the log_2_FC of k-mers with k=4 of all the analyzed conditions were used for Heatmap representation applying k-means clustering (n clusters=3). Heatmaps graphical representations were depicted using ComplexHeatmap R package (v2.8.0) (https://bioconductor.org/packages/release/bioc/html/ComplexHeatmap.html).

### m^6^A-seq and m^6^A-CLIP

m^6^A-seq was performed on FUS^P525L^ SK-N-BE cells as described by Dominissini et al 2013^52^ with some adjustments. Poly-A selection was performed thanks to the Poly(A)Purist-MAG kit (Invitrogen, #AM1922). Poly-A+ RNA was incubated 4min at 94°C for fragmentation and 5*μ*g of fragmented poly-A+ RNA was used for the immunoprecipitation with the anti-m^6^A polyclonal antibody (Abcam, #ab151230). The elution of the immunoprecipitated fraction was carried out thanks to the protocol described in Molinie et al, 2017^53^. m^6^A-CLIP for qRT-PCR validation of the m^6^A-seq was performed as described in Di Timoteo et al.^54^

### MeRIP-Seq analysis

RNA libraries for all samples were produced using Stranded Total RNA Prep with Ribo-Zero Plus (Illumina). All samples were sequenced on an Illumina Novaseq 6000 Sequencing system with an average of about 74million 100nucleotides long paired-end read pairs. Trimmomatic (v0.39)^46^ and Cutadapt (v3.2)^47^ were used to remove adapter sequences and poor quality bases; minimum read length after trimming was set to 35. Using Bowtie2 software (v2.4.2)^55^ reads aligned to contaminant sequences of human rRNAs retrieved from NCBI were discarded. STAR software (v2.7.7a)^48^ was used to align reads to GRCh38 genome using ENCODE standard parameters referred in the manual. PCR duplicates were removed from all samples using MarkDuplicates command from Picard suite (v2.24.1) (https://broadinstitute.github.io/picard/) and multi-mapped and un-properly paired reads were filtered out using bamtools (v2.5.1)^56^ and samtools (v1.7)^57^ respectively. Peak calling was performed using exomePeak2 (v1.9.1) (https://github.com/ZW-xjtlu/exomePeak2) software filtering out peaks with log_2_FC (IP /INP) < 1; adjusted p-value > 0.01 and with less than a total of 10 fragments in input or m^6^A-IP samples. Peaks were annotated with Ensembl reference gtf (release 99) using bedtools (v2.29.1) with *–s –split –wao* parameters. After peak annotation, isoform and gene expression was assessed using RSEM (v1.3.1)^58^ on input samples. Peaks overlapping genes expressed <1FPKM in the INP condition were filtered out. Meta-gene peak coverage and motif enrichment analysis was performed with RNAmod^59^.

Reads coverage for the forward and reverse strand were generated using deepTools bamCoverage software (v3.5.1; https://deeptools.readthedocs.io/en/develop/content/tools/bamCoverage.html).

---

**Fig. S1**

**a**, Densitometry analyses showing the relative protein level of FUS in SK-N-BE cells upon doxyciclin induction (D+) with respect to the control condition (D-). The relative protein quantity in the bars is represented as mean of replicates with standard deviation. Dots represent independent replicates. n=3 biologically independent replicates. The *ratio* of each sample versus its experimental control was tested by two-tailed Student’s t test. P-values are indicated. **b**, Representative images of the indicated SK-N-BE cells either stressed (Stress) or in control (Ctrl) condition. G3BP1 antibody staining is illustrated in green, FUS in red. The nuclei are stained with DAPI (blue). The scale bar is 20*μ*m. **c**, Representative image of a SK-N-BE FUSP525L cell in stress condition (left) used for the signal profile analyses of ten granules (right). The nucleus is stained with DAPI (blue). G3BP1 antibody staining, and the corresponding signal is illustrated in green, FUSP525L in red. The scale bar is 5*μ*m. n=10. **d**, Representative western blots (left) and the corresponding densitometry analyses (right) evaluating the decrease of METTL3 protein upon shRNAs (sh) transfection in SK-N-BE FUS^wt^ or SK-N-BE FUS^P525L^ cell lines; ACT-*β* was used as loading control. Relative protein levels were represented as relative quantities with respect to wild-type cells set to 1. The relative protein quantity in the bars is represented as mean of replicates with standard deviation. Dots represent independent replicates. n=3 biologically independent replicates. The *ratio* of each sample versus its experimental control was tested by two-tailed Student’s t test. P-values are indicated. **e**, Relative quantification of m^6^A on RNA upon METTL3 knock-down (sh) with respect to a control condition (scr, set as 100%) in either FUS^wt^ or FUS^P525L^ SK-N-BE cells. The relative m^6^A percentage in the bars is represented as mean of replicates with standard deviation. Dots represent independent replicates. n=3 biologically independent replicates. The *ratio* of each sample versus its experimental control was tested by two-tailed Student’s t test. P-values are indicated. **f**, Bar plot depicting SG enrichment (log_2_FC SG/INP) of selected RNAs to confirm proper SG purification in D^-^ (black bar), FUS^WT^ (grey bar) and FUS^P525L^ (white bar) conditions: HUWE1, TRIO and AHNAK were used as positive controls while ATP5O as negative one. The log2FC thresholds to define SG enriched RNAs (log2FC > 1) and depleted ones (log2FC < -1) are indicated as dotted grey lines. **g**, Scatter plot depicting RNA differential enrichment in SG in FUS^P525L^ vs FUS^P525L^ METTL3-KD conditions without FUS^P525L^ doxycycline induction. Axes describes log_2_FC of SG RNA enrichment in the indicated conditions. Red dots indicate DMETTL3-enriched RNAs. Blue dots indicate DMETTL3-depleted RNAs. Black dots indicate commonly enriched RNAs that are enriched in SG in both conditions. Grey dots indicate invariant RNAs. n=2. **h**, Venn diagram depicting the overlap between ΔMETTL3 Relocated RNAs and SG enriched RNAs in FUS^WT^ and FUS^P525L^ conditions. **i**, Venn diagrams depicting the overlap among the SG enriched RNAs in the three analyzed condition (D-, FUS^WT^ and FUS ^P525L^) in control cells (left panel) and upon METTL3 downregulation (METTL3 KD, right panel). **j**, Venn diagrams depicting the overlap among the top 5000 (left) and top 500 RNAs (right) ranked by SG enrichment in each analyzed condition (D-, FUS^WT^ and FUS ^P525L^ upon METTL3 depletion). The bar plot (middle panel) displays that there is no strong divergence of FUS^P525L^ SG enriched RNAs comparing to the other conditions (D-, FUS^WT^) in every selected group. **k**, Bar plot depicting the library sizes of RNA-Seq samples before the resampling procedure (upper panel) and after the resampling procedure (lower panel). Library sizes of FUS^P525L^ samples were used as reference in order to resample fragments of other samples. Venn diagram (right panel) displays that after the resampling procedure there is an high overlap between SG enriched RNAs in the three analyzed condition D-, FUS^WT^ and FUS ^P525L^ upon METTL3 depletion. **l**, Heatmap (lower panel) showing k-mers (4-mers) enrichment in SG among different condition. Rows represent each possible k-mer with k=4, while columns represent conditions. K-mers enrichment is described as ratio between mean k-mer percentage in SG enriched RNAs vs invariant RNAs. The three clusters identified using K-means clustering (n clusters=3) indicate k-mers depleted (#1), neither (#2) or enriched (#3) in FUS^P525L^ condition. Color scale describes the magnitude of k-mer enrichment, enrichment values > 1 are represented in red while enrichment values < 1 are represented in blue. Pie charts (upper panel) represents the fraction of GC and AU nucleotides in k-mers clusters. **m**, Sequence logo representing the consensus DRACH motif significantly over-represented in the MeRIP-seq peak regions. **n**, Metagene plot resulted from MeRIP-seq data displaying m^6^A increased signal intensity near the STOP codon. **o**, Representative enrichment of selected RNAs resulted methylated according to MeRIP-seq data. WTAP and ATP5O were used as positive and negative controls, respectively; immunoprecipitation with IgG was used as control. n=2. **p**, Box plots displaying RNA length (Kb) of the indicated RNA groups identified by SG differential enrichment analysis among FUS^P525L^ vs FUS^P525L^ METTL3 KD. Statistical significance was evaluated with Mann-Whitney U test. P-values are indicated in the figure.

**Fig. S2**

**a**, Schematic representation of the METTL3 *locus* and the plasmid donor used for CRISPR/Cas-9 mediated genome editing to insert a degron tag (mAID) at the N-terminus of METTL3. Primers for genotyping are indicated with arrows. The size of the expected amplicon is specified. **b**, Genomic analyses of selected SK-N-BE clones after CRISPR/Cas9 mediated genome editing to insert a degron tag (mAID) at the N-terminus of METTL3. Heterozygotes are red squared. Blue arrows indicate the size of the amplicons deriving from either wild type (METTL3 WT) or edited genome (mAID METTL3). The donor DNA and gDNA from unedited cells were used as positive or negative controls, respectively. **c**, Representative western blot analysis showing METTL3 and METTL14 decrease in the indicated SK-N-BE cells after degron tag insertion (mAID). ACT-β was used as loading control. **d**, Relative protein levels of METTL3 (n=4 biologically independent replicates) and METTL14 (n=3 biologically independent replicates) in the indicated SK-N-BE degron tag (mAID) insertion. Levels were normalized over ACT-β protein level and expressed as relative quantities with respect to wild-type cells set to 1. The relative protein quantity in the bars is represented as mean of replicates with standard deviation. Dots represent biologically independent replicates. The *ratio* of each sample versus its experimental control was tested by two-tailed Student’s t test. P-values are indicated. **e**, Bar plot showing METTL3 relative RNA level decrease in the indicated SK-N-BE cells. Values are normalized against ATP5O transcript and expressed as relative quantity with respect to non-edited cells set to a value of 1. The relative RNA quantity in the bars is represented as mean of the fold change with standard deviation. Dots represent each replicate. n=4 biologically independent replicates. The *ratio* of each sample versus its experimental control was tested by two-tailed Student’s t test. P-values are indicated. **f**, Bar plot showing the m^6^A level decrease in the indicated SK-N-BE cells. Levels are represented as percentage with respect to control cells set as 100%. Each dot represents a single replicate. n=3 biologically independent replicates. The *ratio* of each sample versus its experimental control was tested by two-tailed Student’s t test. P-values are indicated. **g**, Representative western blot analysis showing stable FUS expression in the indicated SK-N-BE cells. GAPDH was used as loading control. **h**, Levels of selected RNAs in a representative stress granules immunoprecipitation in SK-N-BE FUS^WT^ or FUS^P525L^ cell lines, induced for the expression of FUS (D+), upon 1hr stress treatment (A+). RNA levels are expressed as percentage of input normalized on an invariant RNA enrichment (CALM1). ATP5O transcript was used as negative control. The relative RNA enrichment in the bars is represented as mean of the replicates with standard deviation. Dots represent each replicate. n=2 biologically independent replicates. The *ratio* of each sample versus its experimental control was tested by two-tailed Student’s t test. P-values are indicated. **i**, Box plots showing granule volumes in the indicated SK-N-BE cells. Black dots represent the average volumes. Gray dots represent the volume of each measured granule. n=3 biologically independent replicates. Seven fields were acquired for each biological replicate. **j**, Box plots showing number of small, medium, or large granules per cell the indicated SK-N-BE cells. Seven fields were acquired for each biological replicate. n=3 biologically independent replicates. P-values are indicated. **k**, Genomic analyses of selected iPS clones after CRISPR/Cas9 mediated genome editing to insert a degron tag (mAID) at the N-terminus of METTL3. Homozygotes are red squared. Blue arrows indicate the size of the amplicons deriving from either wild type (METTL3 WT) or edited genome (mAID METTL3). The donor DNA and gDNA from unedited cells were used as positive or negative controls, respectively. **l**, Representative western blot analysis (left) showing METTL3 and METTL14 decrease in the indicated SK-N-BE cells after degron tag insertion (mAID). ACT-β was used as loading control. Relative protein levels (right) of METTL3 (n=4 biologically independent replicates) and METTL14 (n=3 biologically independent replicates) in the indicated SK-N-BE degron tag (mAID) insertion. Levels were normalized over ACT-β protein level and expressed as relative quantities with respect to wild-type cells set to 1. The relative protein level in the bars is represented as mean of the replicates with standard deviation. Dots represent biologically independent replicates. n=3. The *ratio* of each sample versus its experimental control was tested by two-tailed Student’s t test. P-values are indicated. **m**, Representative images of METTL3 immunofluorescence on the indicated MNs (left). METTL3 antibody staining is illustrated in red, TUJ1 in grey. The nuclei are stained with DAPI (blue). The merge of the signals from TUJ1 and DAPI is shown. The scale bar is 10 *μ*m. Box plots representing METTL3 signal intensity analyses (right). Seven fields were acquired. Red dots represent signal average. Each gray dots represents a single cell signal. P-values are indicated. The scale bar is 20 *μ*m. **n**, Bar plot showing the m^6^A level decrease in the indicated iPSC derived MN. Levels are represented as percentage with respect to control cells set as 100%. Each dot represents a single replicate. n=3 biologically independent replicates. The relative m^6^A percentage in the bars is represented as mean of the replicates with standard deviation. The *ratio* of each sample versus its experimental control was tested by two-tailed Student’s t test. P-values are indicated. **o**, Bar plot showing relative RNA levels of selected MN differentiation markers in the indicated MN. Values are normalized against GAPDH transcript and expressed as relative quantity with respect to unedited MN set as 1. The relative RNA quantity in the bars is represented as mean of the fold change with standard deviation. Dots represent each replicate. n=3 biologically independent replicates. The *ratio* of each sample versus its experimental control was tested by two-tailed Student’s t test. P-values are indicated. **p**, Representative images of the indicated stressed MNs. G3BP1 antibody staining is illustrated in green, FUS in red, TUJ1 in grey. The nuclei are stained with DAPI (blue). The merge of the signal is shown. The scale bar is 10 *μ*m. **q**, Box plots showing granule volumes in the indicated MN. Black dots represent the average volumes. Gray dots represent the volume of each measured granule. n=4 biologically independent replicates. Seven fields were acquired for each biological replicate. **r**, Box plots showing number of small, medium, or large granules per cell the indicated MN. Seven fields were acquired for each biological replicate. n=4 biologically independent replicates. P-values are indicated.

**Fig.S3**

**a**, Representative images of the indicated SK-N-BE both in stress and in recovery conditions showing the presence of FUS-only granules. G3BP1 antibody staining is illustrated in green, FUS in red. The nuclei are stained with DAPI (blue). The merge of the signal is shown. The scale bar is 10 *μ*m. **b**, Box plot showing the colocalization analysis of G3BP1 and FUS signals in SK-N-BE FUS^P525L^ cells both in stress and in recovery conditions. Seven fields were acquired for each biological replicate. n=3 biologically independent replicates. The *ratio* of each sample versus its experimental control was tested by two-tailed Student’s t test. P-values are indicated. **c**, Box plot showing the comparison of granule volumes in stress or in recovery conditions. Black dots represent the average volumes. Gray dots represent the volume of each measured granule. Seven fields were acquired for each biological replicate. n=3 (stress) n=4 (recovery) biologically independent replicates. The *ratio* of each sample versus its experimental control was tested by two-tailed Student’s t test. P-values are indicated.

**Fig.S4**

**a**, Bar plot showing the m^6^A level decrease of SK-N-BE FUS^P525L^ cells treated either with DMSO or STM-2457. Levels are represented as percentage with respect to control cells set as 100%. Each dot represents a single replicate. n=3 biologically independent replicates. The *ratio* of each sample versus its experimental control was tested by two-tailed Student’s t test. P-values are indicated. **b**, Box plots representing the number of stress granules per cell in stressed SK-N-BE FUS^P525L^ treated either with DMSO or STM-2457. The red dot indicates the average number of granules in each sample. Each black dot represents the number of granules in a single field. Seven fields were acquired for each biological replicate. n=3 biologically independent replicates. The *ratio* of each sample versus its experimental control was tested by two-tailed Student’s t test. P-values are indicated. **c**, Box plots showing the percentage of unrecovered SK-N-BE FUS^P525L^ treated either with DMSO or STM-2457. The red dot indicates the average percentage in each sample. Each black dot represents the percentage in a single field. Seven fields were acquired for each biological replicate. n=3 biologically independent replicates. The *ratio* of each sample versus its experimental control was tested by two-tailed Student’s t test. P-values are indicated. **d**, Bar plot showing the m^6^A level decrease of iPSC-derived MN FUS^P525L^ treated either with DMSO or STM-2457. Levels are represented as percentage with respect to control cells set as 100%. Each dot represents a single replicate. n=3 biologically independent replicates. The *ratio* of each sample versus its experimental control was tested by two-tailed Student’s t test. P-values are indicated. **e**, Bar plot showing the m^6^A level decrease of patient-derived fibroblasts treated either with DMSO or STM-2457. Levels are represented as percentage with respect to control cells set as 100%. Each dot represents a single replicate. n=3 biologically independent replicates. The *ratio* of each sample versus its experimental control was tested by two-tailed Student’s t test. P-values are indicated. **f, g, h, i, j, k** Box plots representing the S_conf_ or D of FUS^P525L^ or G3BP1 in living SK-N-BE cells treated either with DMSO or STM-2457 under stress or recovery condition. The red dot indicates the average value. Each black dot a single measurement. n=2 biologically independent replicates. The *ratio* of each sample versus its experimental control was tested by two-tailed Student’s t test. P-values are indicated.

