## Supplementary figures for "M^6^A reduction relieves FUS-associated ALS granules"

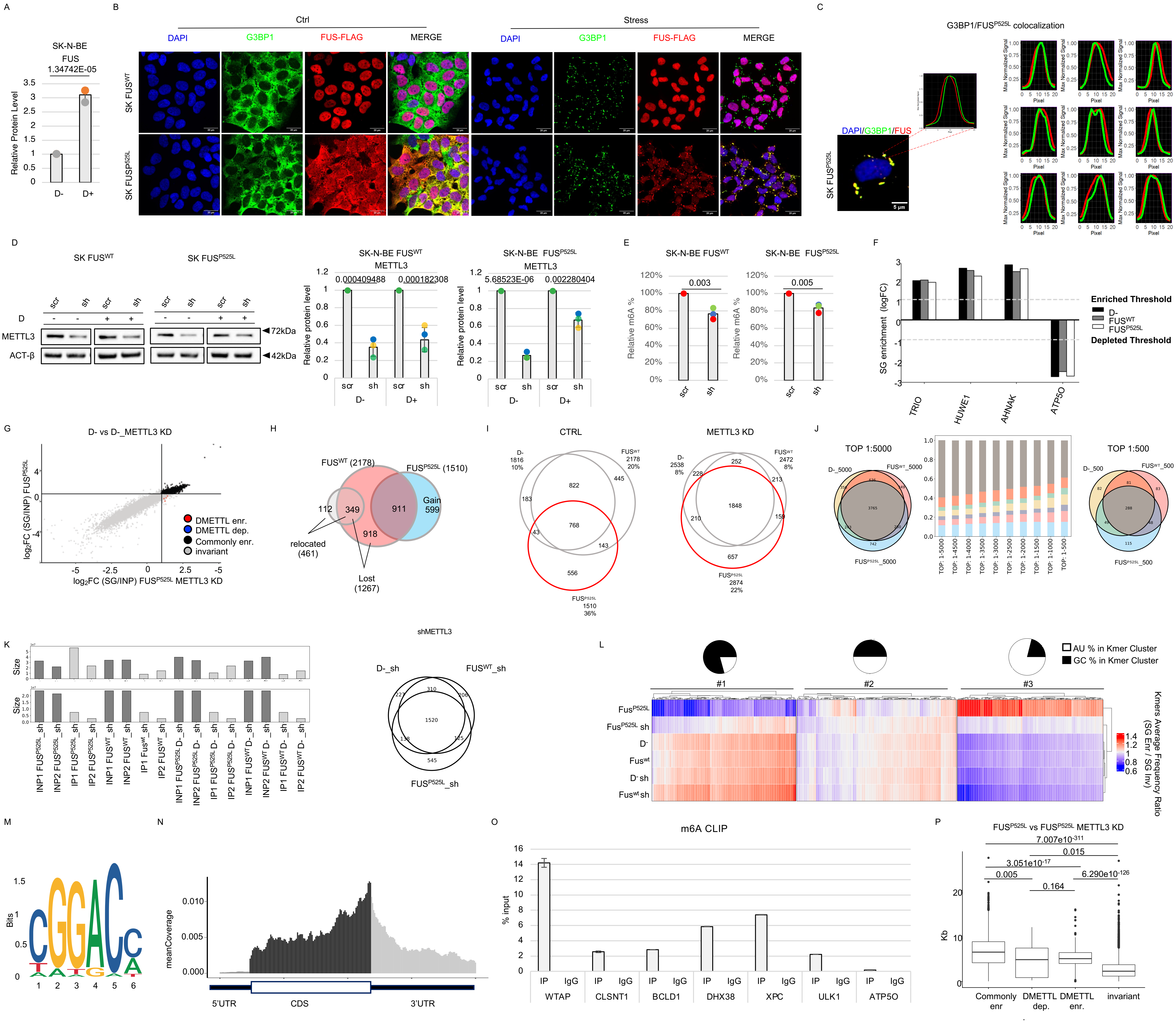


**Fig.S1 METTL3 downregulation restores the physiological RNA composition of stress granules in ALS cellular models**

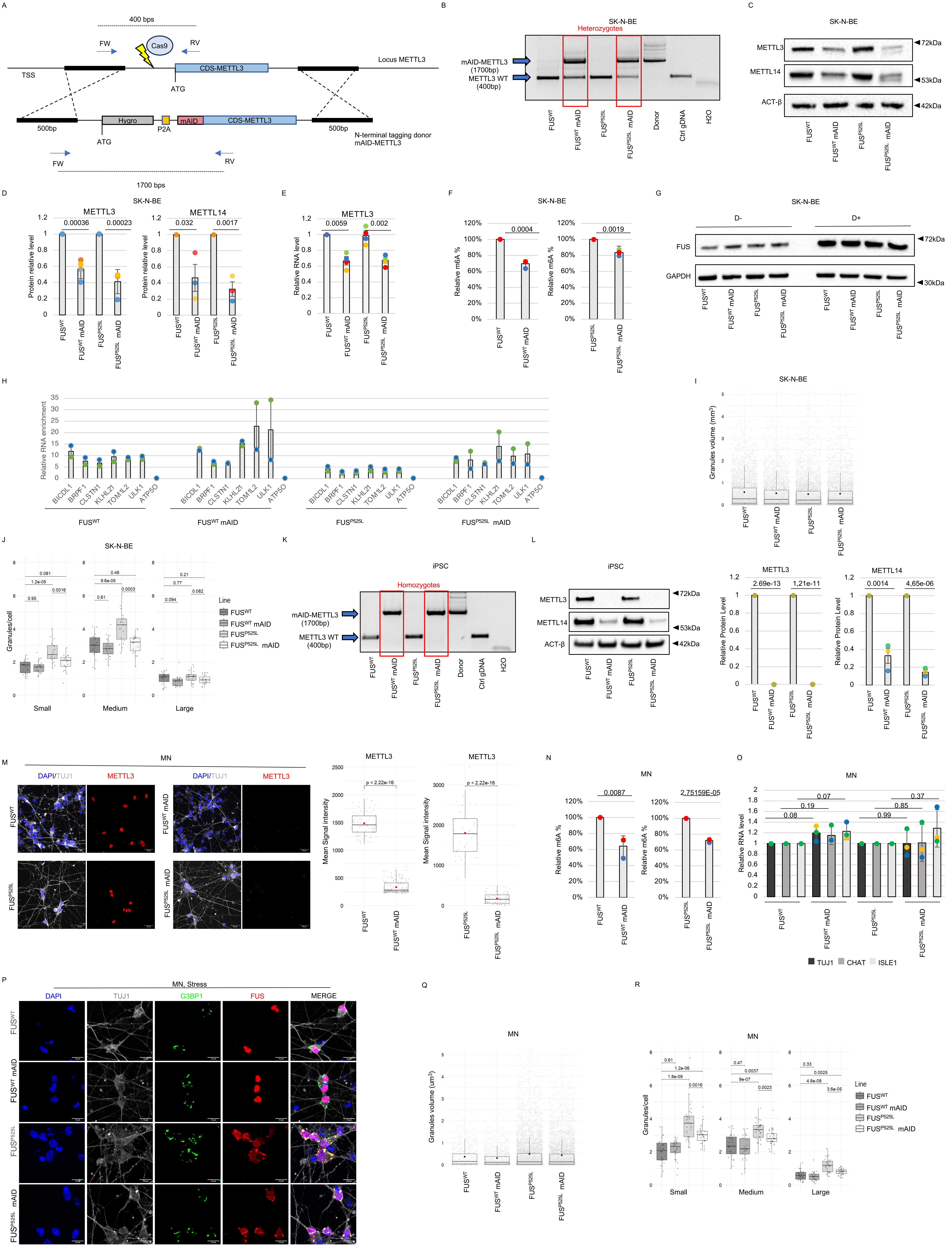


**Fig.S2 METTL3 downregulation reduces the number of SG in ALS cellular models**

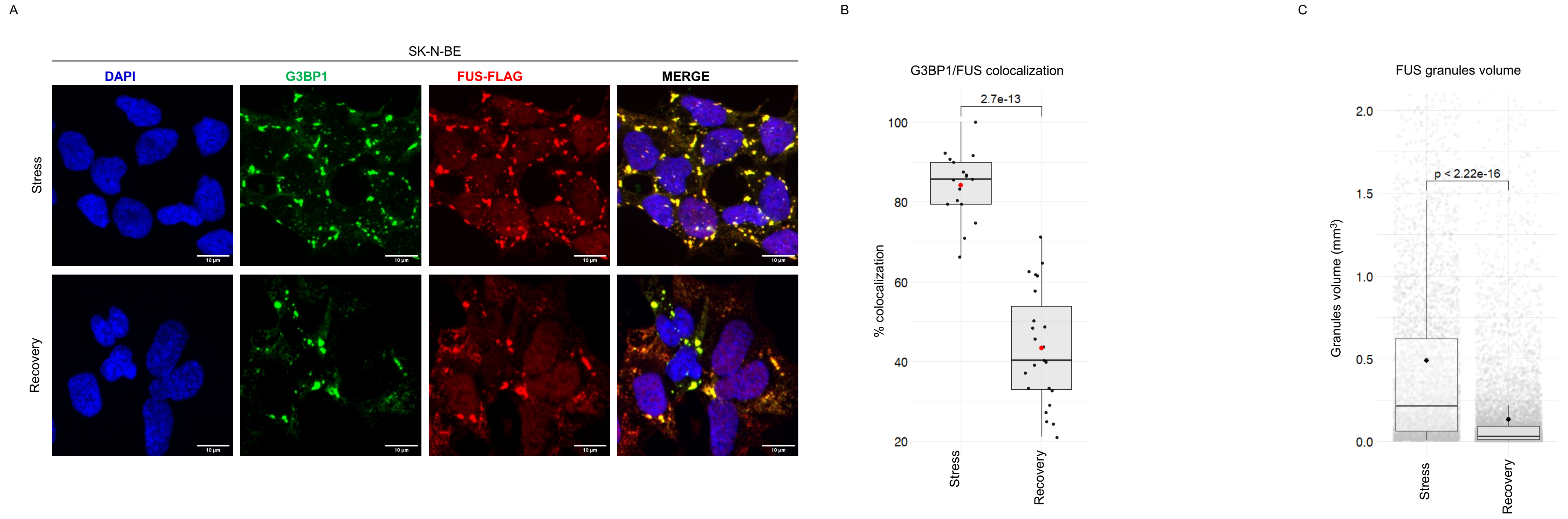


### Fig.S3 METTL3 decrease restores SG recovery rate in ALS cellular models

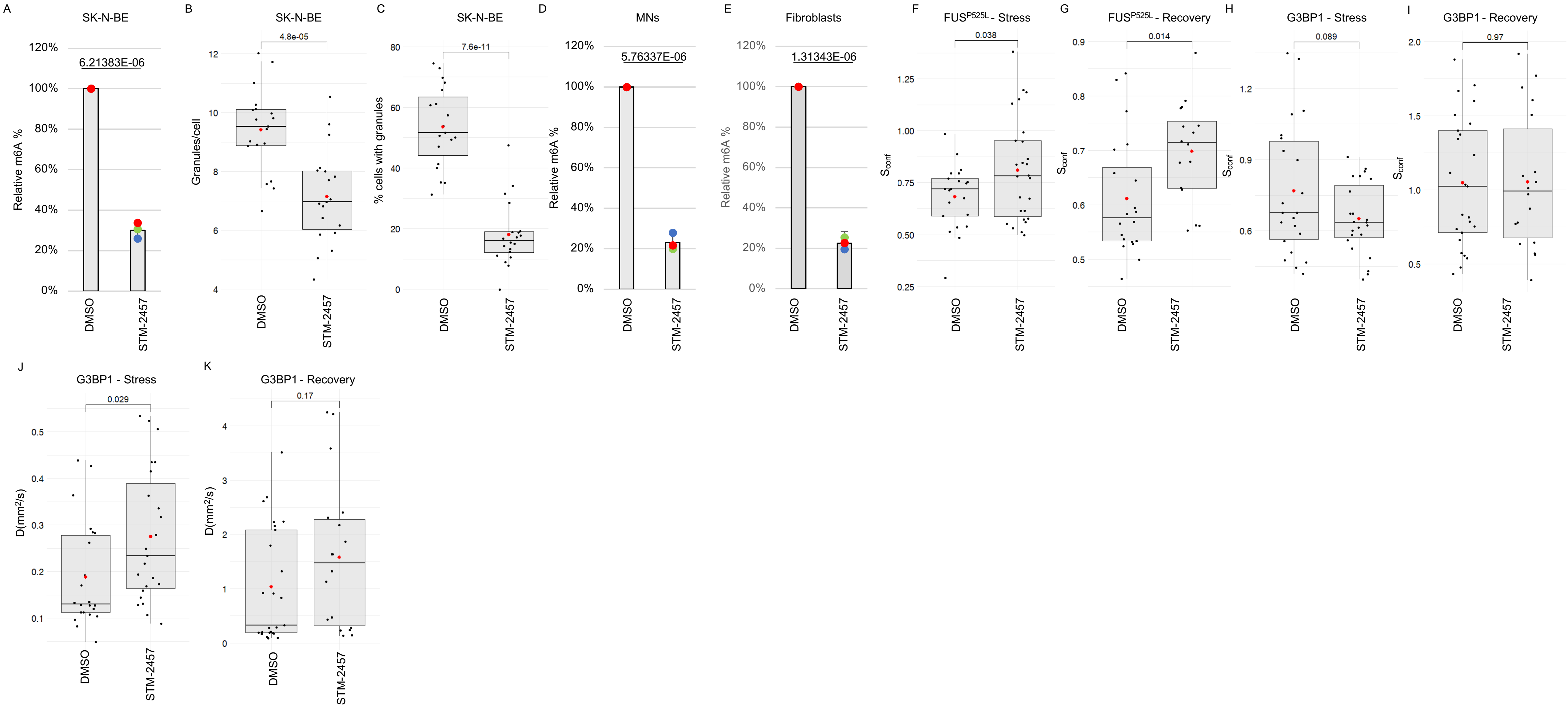


### Fig.S4 METTL3 chemical inhibition relieves FUS-containing SG
